# Assessment of Operant Learning and Memory in Mice Born through Intracytoplasmic Sperm Injection

**DOI:** 10.1101/2020.02.11.942235

**Authors:** M. Lewon, Y. Wang, C. Peters, M. Peterson, H. Zheng, L. Hayes, W. Yan

## Abstract

**Study question:** Are there differences in operant learning and memory between mice born through intracytoplasmic sperm injection (ICSI) and naturally-conceived control (CTL) mice?

**Summary answer:** ICSI females exhibited deficits in acquisition learning relative to CTL females, whereas ICSI males exhibited deficiency in discrimination learning and memory relative to CTL males during initial assessments. ICSI and CTL groups exhibited equally poor long-term retention of learned discrimination and memory performances at old age.

**What is known already:** Some human outcome studies have suggested that ICSI might be associated with an increased risk of certain cognitive disorders, but only one of two behavioral studies with ICSI mouse models have reported differences between ICSI and CTL females. No studies to date have investigated associative learning in ICSI mice.

**Study design, size, duration:** 36 ICSI mice (18 male, 18 female) and 37 CTL mice (19 male, 18 female) aged 3-6 months were compared in a series of operant learning procedures that assessed acquisition of a new behavior, discrimination learning, and memory. 16 ICSI mice (9 male, 7 female) and 17 CTL mice (10 males, 7 females) received follow-up discrimination learning and memory assessments at 12 months of age (six months after the end of initial training) to evaluate retention and reacquisition of learned performances.

**Participants/materials, setting, methods:** Mice received daily operant learning sessions in experimental chambers in which all stimulus events and the recording of responses were automated. Food rewards were delivered for responding under different conditions of reinforcement, which varied by procedure. Subjects received a successive series of sessions of nose poke acquisition training, discrimination training, and the delayed non-matching-to-position (DNTMP) memory procedure. Mixed repeated measures ANOVAs in which the between-subjects factor was group (ICSI vs. CTL) and the within-subjects factor was repeated exposures to learning procedures (i.e., sessions) were used to analyze data.

**Main results and the role of chance:** In comparisons between all mice (i.e., males and females combined), CTL mice exhibited superior performance relative to ICSI in response acquisition (p = 0.03), discrimination (p = 0.001), and memory (p = 0.007). Sex-specific comparisons between the groups yielded evidence of sexual dimorphism. ICSI females exhibited a deficit in acquisition learning relative to CTL females (p < 0.001) but there was not a significant difference between CTL and ICSI males. In the discrimination and memory tasks, ICSI males exhibited deficits relative to CTL males (p = 0.002 and p = 0.02, respectively) but the differences between females in these tasks were not significant. There was no difference in discrimination or memory retention/re-acquisition assessments conducted with mice at 12 months of age. ICSI males and females weighed significantly more than CTL counterparts at all points during the experiment.

**Limitations, reasons for caution:** The study was not blinded. All learning assessments utilized food reward; other assessments of operant, Pavlovian, and nonassociative learning are needed to fully characterize learning in ICSI mice and speculate regarding the implications for cognitive function in humans conceived via ICSI.

**Wider implications of the findings:** Studying learning and memory processes in mouse models has the potential to shed light on ICSI outcomes at the level of cognitive function. Future research should use multiple learning paradigms, assess both males and females, and investigate the effects of variables related to the ICSI procedure. Studying cognitive function in ICSI is an interdisciplinary endeavor and requires coordination between researchers at the genetic and psychological levels of analysis.

**Study funding/competing interest(s):** This work was supported, in part, by grants from NIH (P30GM110767, HD071736 and HD085506 to WY), the Templeton Foundation (Grant ID: 61174 to WY), and a New Scholarly Endeavor Grant from the University of Nevada, Reno Office of Research and Innovation (to ML, YW, HZ, LH, and WY). The authors declare no competing interests.

## Introduction

Intracytoplasmic sperm injection (ICSI) is an assisted reproductive technology (ART) that is achieved through the injection of a single spermatozoon directly into the cytoplasm of an oocyte. ICSI has proven to be effective in treating severe forms of male factor infertility that are difficult to treat with other ARTs. Since the first ICSI pregnancies in 1992 (Palermo et al., 1992), the procedure has grown in popularity and is now the most commonly used ART worldwide (Rozenwaks & Pereira, 2017). In the United States, ICSI use increased from 36.4% of all fertility treatment cycles in 1996 to 76.2% in 2012 (Boulet et al., 2015). Although ICSI was originally developed specifically to treat infertility related to semen quality, the use of ICSI for non-male factor infertility has also increased from 15.4% in 1996 to 66.9% in 2012 (Boulet et al., 2015).

The use of ICSI as the treatment of choice for various types of infertility has raised concerns regarding its overuse, especially in light of the possibility of adverse postnatal outcomes (Esteves et al., 2018). ICSI has been responsible for over two million births since its inception (Palermo et al., 2017). As the earliest ICSI babies are now reaching maturity, researchers have become increasingly concerned with examining ICSI outcomes in various domains. Human outcome studies inherently contain many confounds and biases and therefore must be interpreted with caution (Fauser et al., 2014; Pereira et al., 2017). Nevertheless, some studies have found that ICSI may be associated with increased risks of chromosomal and epigenetic irregularities (Manipalviratn et al., 2009; Odom & Segars, 2010), congenital birth defects (Lacamara et al, 2017; Massaro et al., 2015; Pandey et al., 2012), and cognitive disorders (Hansen et al., 2018; Sandin et al., 2013).

While some studies have suggested a tentative relationship between ICSI and abnormal psychological development, it is particularly difficult to draw conclusions regarding this relationship from human outcome studies because cognitive development is profoundly influenced by individuals’ environmental circumstances (Hart & Risley, 1995; Novak & Peláez, 2004). The heterogeneity of the cultural, familial, and educational environments of children conceived *via* ICSI makes it impossible to extricate the respective contributions of genetic/epigenetic and environmental variables on psychological development. Characterizing the relationship between ICSI and psychological function would ideally involve studying learning and cognitive development in individuals conceived *via* ICSI in well-controlled environments.

This approach is not feasible with humans, but animal models provide an opportunity to control for many environmental factors and study behavior and learning processes that serve as a common basis for cognitive function in humans and nonhumans alike. We were able to identify only two studies that compared ICSI mice to naturally-conceived control (CTL) mice for this purpose. Fernández-Gonzalez et al. (2008) compared ICSI and CTL CD-1 male and female mice in a series of behavioral assays that included an open field test to assess locomotion, an elevated plus maze task to assess sensitivity to anxiety-inducing stimuli, and a free-choice y-maze task to assess habituation to novelty. They found no differences between ICSI and CTL males in any of the procedures, but ICSI females exhibited less exploration in the open field, increased anxiety as measured by time spent in the open arms of the elevated plus maze, and less habituation as measured by time spent in a previously explored arm of the y-maze. Kohda et al. (2011) found no significant differences between male ICSI and CTL C57BL/6 x DBA/2 (BDF1) mice in a series of tests designed to assess locomotion and sensitivity to fear- and pain-inducing stimuli. Female mice were not assessed in the latter study.

The procedures used in the studies cited above allowed for comparisons between ICSI and CTL mice in terms of a) general activity/locomotion, b) sensitivity to anxiety- and pain-inducing aversive stimuli, and c) habituation to novel environmental stimuli. All of these may provide important information relevant to psychological function, but only the procedures that measured habituation (Fernández-Gonzalez et al., 2008) may be considered to assess *learning* per se. Learning is defined generally as changes in organisms’ behavior with respect to particular environmental events or stimuli as a result of previous experiences (Pierce & Cheney, 2013). Habituation is one of the most basic learning processes and describes situations in which an animal’s response to a particular environmental stimulus or event decreases with repeated exposure to that stimulus or event (Groves & Thompson, 1970; Rankin et al., 2009; Thompson & Spencer, 1966). Habituation is categorized as an example of *nonassociative learning* because changes in behavior occur simply through exposure to an environmental stimulus (Domjan, 2015).

*Associative learning* is a higher form of learning and serves as the basis for cognition in all organisms, including humans (Domjan, 2015; Ginsburg & Jablonka, 2010; Mackintosh, 1974). There are two fundamental associative learning processes that have been studied extensively with both humans and nonhumans since the early 1900s: Pavlovian learning (Domjan, 2005; Pavlov, 1927/1960; Rescorla, 1988) and operant learning (Pierce & Cheney, 2013; Skinner, 1938, 1953; Thorndike, 1911). In Pavlovian learning, organisms learn about relations between environmental stimuli. If two stimuli frequently occur together in organisms’ environments, they come to respond to the two stimuli in a similar fashion. This allows organisms to prepare for and more effectively interact with biologically important stimuli (Domjan, 2005). In operant learning, organisms learn about relations between their behavior and its effects on the environment. Responses that regularly produce rewarding consequences (e.g., the opportunity to eat food, drink water, or escape from aversive stimuli) will come to occur more frequently in the environmental settings where they have been associated with these consequences. Responses that do not produce rewarding consequences, or result in exposure to aversive events, come to occur less frequently. Pavlovian and operant learning allow organisms to interact with their environments effectively and adapt to changes in the environment that occur during their lifetimes. These learning processes serve as the basis for language and other forms of complex human behavior (De Houwer et al., 2016; Jablonka & Lamb, 2014; Sturdy & Nicoladis, 2017).

To date, there have been no studies that compared associative learning between ICSI and CTL mice. Studying these fundamental learning processes has the potential to provide insights into relationships between ICSI and cognitive function that may not be obtained from human outcome studies. The purpose of the present study was to conduct the first assessment of operant learning and memory in a mouse model of ICSI. ICSI and naturally-conceived CTL mice were exposed to a series of operant learning procedures that assessed acquisition of a new behavior, discrimination learning, and memory. These assessments were conducted while the mice were between 3-6 months of age. Follow-up assessments were then conducted with some of the mice to investigate retention and re-acquisition of learned performances when the mice were 12 months of age.

## Methods and Materials

### Naturally-Conceived Control (CTL) Mice

All animal work was performed following the protocol approved by the Institutional Animal Care and Use Committee (IACUC) of the University of Nevada, Reno. Adult (6-8 weeks of age) CD-1 mice used in this study were purchased from Charles River, and housed under pathogen-free conditions in a temperature- and humidity-controlled animal facility at the University of Nevada, Reno. Natural mating was set up by placing one adult male into a cage with one adult female, and all of the naturally-conceived control (CTL) mice used in this study were those from the first 4 litters of four breeding pairs. Pups were weaned at 3 weeks after birth.

### Intracytoplasmic Sperm Injection (ICSI) Mice

Adult female CD-1 mice at 6-12 weeks of age with body weight ranging between 25-45 grams were used as either egg donors or recipients/surrogates. These female mice were superovulated by intraperitoneal injection of 7 IU of Pregnant Mare’s Serum Gonadotropin (PMSG), followed by intraperitoneal injection of 7 IU of human Chorionic Gonadotropin (hCG) 48 h later. Mature oocytes (MII stage) were collected from the oviducts 14-16 h after hCG injection, and freed from cumulus cells by treatment with 1.5mg/ml bovine testicular hyaluronidase (Sigma, Cat# H3506) in the M2 medium (Millipore, Cat# MR-015-D) at 37°C for 2 min. The cumulus-free oocytes were washed and kept in the KSOM+AA medium (Millipore, Cat#MR-121-D) in an incubator (Sanyo, Cat# 19AIC) at 37°C with air containing 5% CO_2_ before ICSI.

ICSI was performed as described previously (Stein and Schultz 2010; Yuan, et al. 2015), with minor modifications. In brief, WT cauda epididymal sperm were collected into 1 ml HTF medium (Millipore, Cat# MR-070-D), followed by incubation for ∼30 min at 37°C in an incubator with humidified air containing 5% CO_2_, allowing spermatozoa to swim into the medium. The top 100 µl sperm suspension was sonicated at the medium level for five times with 3 seconds each (Bioruptor UCD-200; Diagenode). An aliquot of 2 µl sperm HTF suspension was mixed immediately with 50 µl of 4% PVP (Sigma, Cat# P5288) in water (Millipore, Cat# TMS-006-C). A single sperm head was picked up and injected into the mature oocytes using a glass pipette equipped with a piezo drill under the control of an electric micromanipulator (TransferMan NK2, Eppendorf). Injection of ∼20 oocytes was completed within 20 minutes at room temperature. Sperm sonication was then repeated to obtain freshly prepared sperm heads for injection. Injected oocytes were transferred to the KSOM+AA medium (Millipore, Cat# MR-121-D) covered by mineral oil and cultured in an incubator at 37°C with humidified air containing 5% CO_2_. Between 4-6 h post ICSI, 18-26 2PN stage embryos were transferred into the oviducts of pseudo-pregnant CD-1 females (8-16 weeks of age) that had been mated during the prior night with vasectomized adult CD-1 males (10-16 weeks of age).

### Subjects

36 ICSI (18 males and 18 females) and 37 naturally-conceived CTL mice (19 males and 18 females) obtained as described above served as the subjects. All the mice were between 12-13 weeks of age at the beginning of the training described below.

### Housing

ICSI and CTL mice were housed separately in clear plastic Tecniplast® home cages in same-sex groups of three to five mice per cage. Cages were equipped with absorbent corn cob bedding and items for enrichment including cotton fiber nestlets, a transparent red polycarbonate mouse hut and wooden gnawing sticks. Cages were housed in a temperature- and humidity-controlled colony room with a 12:12 light/dark cycle with lights on at 7:00. Except for the scheduled deprivations, subjects had free access to laboratory chow (Harlan Teklad) in overhead feeders. Subjects had free access to purified drinking water at all times.

### Food Deprivation

In order to establish motivation for the sucrose pellet rewards used in experimental sessions, subjects were deprived of food 14 h prior to daily experimental sessions. Food was removed from the subjects’ cages daily at 19:00. Mice had free access to water during the food deprivation period. Experimental sessions were conducted daily at 9:00, and food was returned to the cages after all mice had completed their training sessions. They then had free access to food and water until the next deprivation period.

### Handling and Weighing

Mice were handled using 15 cm tall x 5.75 cm diameter clear plastic tubes open on one end and wide enough to allow the subjects to move freely while sitting in the bottom. Handling tubes have been shown to reduce inter-handler variability and handler-induced stress (Hurst & West, 2010). Prior to each session, a mouse was guided into the tube, weighed, and then placed in the experimental apparatus. When the session concluded, the mouse was transported back to its home cage in the tube.

### Apparatus

All learning and memory assessments were conducted in Med Associates® (St. Albans, VT) modular operant test chambers (ENV-307A). The inside dimensions of the chambers were 12.7 cm high x 15.9 cm wide x 14.0 cm deep. Side walls were composed of transparent polycarbonate, and the front and back walls were composed of three modular columns of aluminum panels. Each chamber was housed in a sound attenuating cabinet with a ventilation fan to mask ambient noise. A 100 mA house light (ENV-315M) was mounted in the center column of the back wall of the chambers 10 cm above the grid floor. On the front wall of the chambers, opposite of the house light, a receptacle measuring 3.8 cm high x 8.9 cm wide was mounted in the center column 0.5 cm above the grid floor. The receptacle was capable of receiving 20 mg Bio-Serv sucrose reward pellets delivered via a pedestal mount pellet dispenser (ENV-203M-20). Two illuminable nose poke operanda (ENV-313M) were mounted 3 cm to either side of the receptacle. The access port for each nose poke measured 1.3 cm in diameter x 1 cm deep. Entry of a subjects’ nose at least 0.64 cm into the access port broke a photobeam and defined a response. The presentation and recording of all experimental events were controlled via MED-PC IV (Med Associates) software.

### Magazine Training

Prior to the learning and memory assessments described below, magazine training was provided to teach the subjects to approach the food receptacle and eat when reward pellets were delivered. Subjects were 12-13 weeks of age at the onset of this training and were deprived of food prior to all sessions as described above. Once an animal was placed inside the chamber, a single pellet was delivered when the animal was oriented toward the receptacle but did not have its head inside of it. After the animal approached and ate the pellet, another pellet was delivered in the same manner. A session was terminated when a mouse had consumed seven pellets. The latency between the delivery of a pellet and its consumption was recorded for each pellet. Each mouse received two such sessions per day for five consecutive days (10 total sessions). By the end of this training, all subjects reliably approached the receptacle and consumed pellets when they were delivered.

### Learning and Memory Assessments

Subjects were exposed to four operant learning and memory assessments conducted in succession. These procedures were the same as those described in Lewon et al. (2017). Each successive assessment was designed to evaluate an increasingly complex performance. These are described below.

### Nose Poke Acquisition

The first assessment was designed to evaluate the acquisition of a new response through reinforcement. Reinforcement describes a fundamental learning process whereby the frequency of a behavior increases because it has been followed by a rewarding consequence (Domjan, 2015). In the present study, the behavior to be acquired was nose poking (i.e., insertion of the nose at least 0.64 cm into the portal of the nose poke operanda) and the rewarding consequence was the delivery of a sugar pellet. The frequency with which this behavior increased through reinforcement and occurred across training sessions provided a measure of acquisition learning.

Subjects were 12.5-13.5 weeks of age at the beginning of this assessment. Each session began with the illumination of the house light and both nose poke stimulus lights. Responses on either nose poke were immediately followed by the delivery of one sucrose pellet (i.e., a fixed-ratio 1 schedule of reinforcement). Each session was terminated after 15 minutes. One session was conducted daily across 10 consecutive days.

### Switching Discrimination Task

The purpose of the second procedure was to assess discrimination learning. Discrimination occurs when organisms learn to engage in a response when the probability of reinforcement is high while abstaining from responding when the probability of reinforcement is low. Discrimination learning tasks may take many forms, but the most common procedure involves rewarding a response when it occurs in one environmental context but withholding reward when the response occurs in a different context. Evidence of discrimination learning is obtained when the response comes to occur more frequently in the setting where it is rewarded and less frequently in settings where it is not. Discrimination learning serves as the basis for many activities that are considered to be cognitive in nature, and abnormalities in this domain are characteristic of a wide range of psychological disorders (Domjan, 2015).

We assessed discrimination learning in a series of sessions in which responses that occurred on illuminated nose pokes were rewarded while responses that occurred on unilluminated nose pokes were not. All mice were 14-15 weeks of age at the beginning of this training. Each session began with the illumination of the house light and the start of a trial in which one of the two nose pokes was illuminated (the program arranged it such that there was a 0.5 probability of either). Responses on the unilluminated nose poke were recorded but produced no programmed consequences. A response on the illuminated nose poke was rewarded with the immediate delivery of a sugar pellet followed by a 5-s intertrial interval (ITI) before the commencement of the next trial. Because there was a 0.5 probability of either nose poke being illuminated on any given trial, the subjects were required to learn to respond on the illuminated nose poke, regardless of position (thus the name *switching discrimination task*; SDT). Sessions were terminated after 15 minutes, and one session was conducted daily for 20 consecutive days.

Discrimination index (DI) provided a measure of the extent to which this discrimination performance was learned. DI was calculated by dividing the total number of responses on the illuminated nose pokes by the total number of responses on the illuminated and unilluminated nose pokes during a session. As we have noted, evidence of discrimination learning is provided by higher response frequencies in settings in which responses have been reinforced (i.e., illuminated nose pokes) relative to settings in which they have not been reinforced (i.e., unilluminated nose pokes). Higher DI values therefore represent greater discrimination learning.

### Delayed Non-Matching-To-Position Memory Task

This task was designed to assess memory. The delayed non-matching-to-position procedure (DNMTP; Steckler et al., 1998) was chosen because it is held to assess two types of memory: working memory and reference memory. Memory researchers describe working memory as information that is retained only long enough to complete a particular task immediately at hand. Once the task is completed, the information is no longer necessary/relevant. On the other hand, reference memory refers to the longer-term retention of information that allows for the successful use of shorter-term working memory in the completion of a task. According to memory theorists, reference memory provides the context necessary to appropriately use working memory (Domjan, 2015).

The DNMTP procedure proceeded as follows. Each session began with the illumination of the house light and the start of a trial in which one of the two nose pokes was illuminated (0.5 probability of either). This portion of the trial was called the forced choice portion: mice were required to respond on the illuminated nose poke to proceed to the subsequent portions of the trial. If they responded on the unilluminated nose poke, there were no programmed consequences. A response on the illuminated nose poke initiated a 2-s retention interval during which both nose pokes were dark Any responses that occurred during this interval produced no programmed consequences. Following the retention interval, both nose pokes were illuminated for the free choice portion of the trial, and subjects could respond on either nose poke. Responses on the same nose poke as required during the forced choice portion of the trial were counted as incorrect and no reward was delivered. Responses on the opposite nose poke of the forced choice trial were counted as correct and rewarded with the delivery of a sugar pellet (thus the name *non*-*matching-to-position*). A trial ended after a correct or incorrect response on the free choice portion and was followed a 5-s ITI. After the ITI, the next trial began with another forced choice. Sessions were terminated when an animal completed 20 trials or 30 minutes, whichever occurred first. Subjects were 17-18 weeks of age at the beginning of this training and received one session daily for 30 consecutive days.

In order to obtain rewards in a trial, mice were required to respond on the nose poke that was not the one on which they responded in the forced choice portion. The working memory aspect of this performance was that the mice had to remember where they had responded in the forced choice portion of the trial during the retention interval. The reference memory portion involved remembering the general rule for reward: respond on the nose poke opposite of the one on which they responded during the forced choice portion of the trial, whether it occurred on the left or right nose poke. When the mice did so, they received a sugar pellet reward and the trial was counted as “correct.” The proportion of correct trials per session provided a measure of memory performance.

### DNMTP Retention Checks

After the 30 trials of DNMTP training described above, mice were removed from the training environment for a prescribed period of time before receiving three additional DNMTP retention check sessions to assess long-term memory of the DNMTP performance. Sessions were identical to those described above. The first retention check occurred two days after the last DNMTP training session. The second occurred five days after the first, and the third occurred 10 days after the second. Subjects were between 21-23 weeks of age during the three retention checks.

### Follow-Up Assessments with Aged Mice

After the initial battery of assessments, follow-up assessments were conducted with some of the same mice from the initial assessments (CTL n = 17; 10 males, 7 females; ICSI n = 16, 9 males, 7 females) when they were between 52-53 weeks of age (i.e., approximately 30 weeks after the last DNMTP retention check session). Prior to the follow-up assessments, mice were weighed for five days under free-feeding conditions starting at 52 weeks of age. After five days, the food deprivation schedule described above was imposed and assessments commenced. The follow-up assessments consisted of 15 daily sessions of the switching discrimination task followed immediately by 15 daily sessions of the DNMTP memory task. All subjects had previous exposure to these procedures during their initial training, and the follow-up assessments were therefore designed to test retention and re-acquisition of these performances at old age.

### Statistical Analysis

Mixed repeated measures ANOVAs were used to compare the results for ICSI and CTL mice in each learning and memory assessment. The between-subjects factor in these analyses was group (ICSI vs. CTL) and the within-subjects factor was session. Omnibus analyses were used to compare all ICSI and CTL mice, and these were followed by sex-specific analyses (i.e., ICSI vs. CTL males and ICSI vs. CTL females). The analyses tested for main effects of group and session as well as for a group x session interaction. We used an *α* value of 0.05 as the criterion for significance, and partial-eta squared values (ηp^2^) are provided as estimates of effect sizes.

## Results

### Nose Poke Acquisition

The training sessions in this phase of the experiment were designed to assess the acquisition of a new behavior through reinforcement learning. Figure 1 shows the mean number of responses per session for all ICSI and CTL subjects of both sexes (top) and separated by ICSI and CTL males and females (bottom). The top panel shows that subjects in both groups generally made more responses in each subsequent training session, but the CTL subjects made slightly more responses in all but one session. The bottom panels show no consistent differences between ICSI and CTL males, but CTL females consistently made more responses per session than their ICSI counterparts. This means that the slight overall difference between all ICSI and CTL mice shown in the top panel of Figure 1 is largely due to rather large and consistent differences in the number of responses per session between ICSI and CTL females during this procedure.

**Figure 1.**
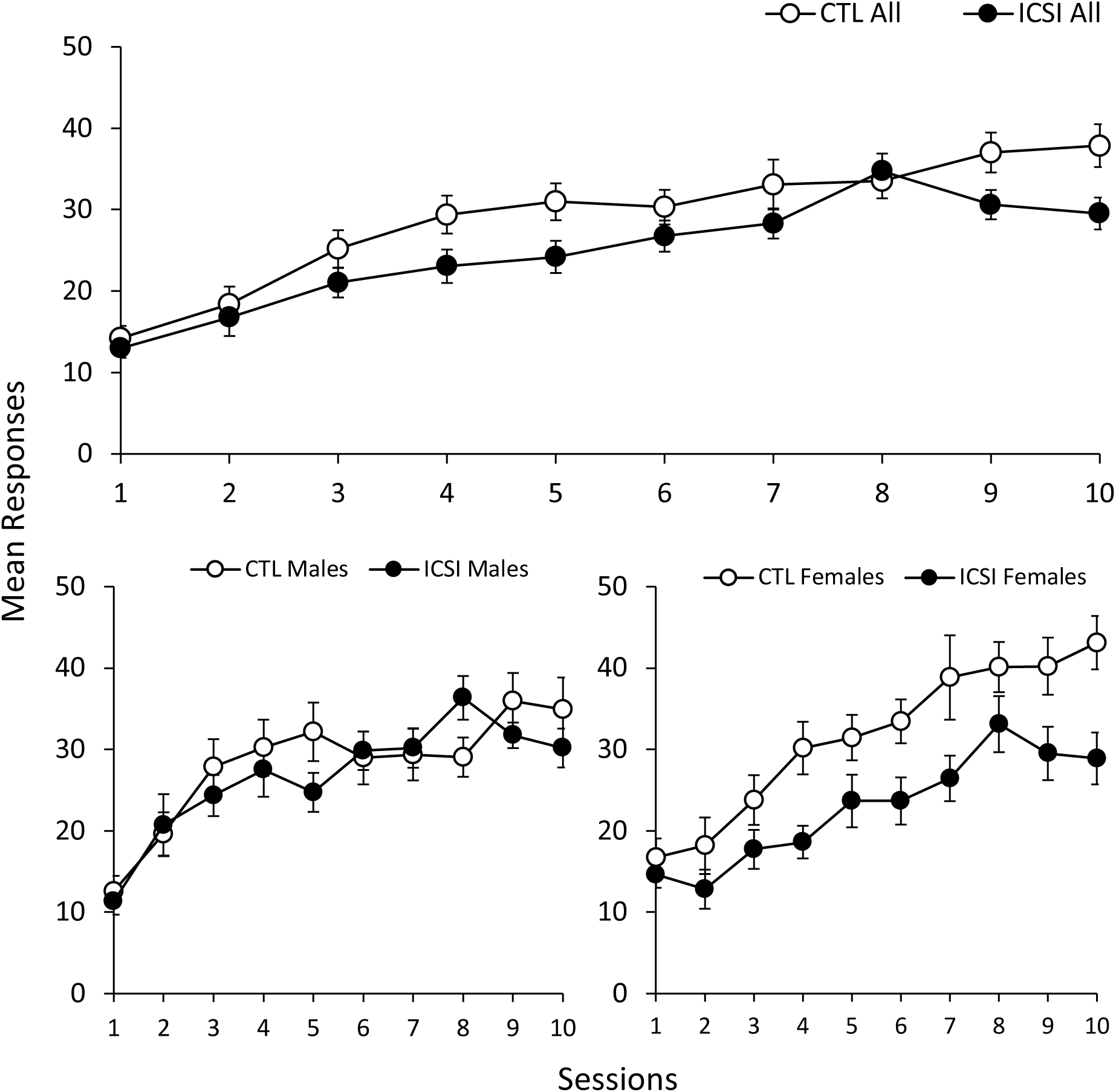
Mean responses per session (+/- standard error of the mean, SEM) during nose poke acquisition sessions for all ICSI and CTL mice (top) and separated by ICSI and CTL males and females (bottom).

The mixed repeated measures ANOVA comparing all ICSI and CTL mice (males and females combined) found a large effect for session (F_9, 639_ = 44.31, p < 0.001, ηp^2^ = 0.38) and smaller effects for group (F_1, 71_ = 4.93, p = 0.03, ηp^2^ = 0.07) and the group x session interaction (F_9, 639_ = 2.06, p = 0.03, ηp^2^ = 0.03). The same analysis was used to compare ICSI and CTL males and found a large effect for session (F_9, 315_ = 20.54, p < 0.001, ηp^2^ = 0.37). There was a barely significant effect for the group x session interaction (F_9, 315_ = 1.96, p = 0.05, ηp^2^ = 0.05), but there was no main effect for group. The comparison between ICSI and CTL females found significant main effects for session (F_9, 306_ = 28.46, p < 0.001, ηp^2^ = 0.46) and group (F_1, 34_ = 6.98, p = 0.01, ηp^2^ = 0.17) but no significant group x session interaction.

To summarize, there was little difference in acquisition between ICSI and CTL males, but the CTL females acquired nose poke responding more readily than the ICSI females. While the CTL females consistently made more responses per session than ICSI females, the statistical analysis did not find a significant group x session interaction. It appeared that CTL females consistently responded more than ICSI females, but the degree to which responding increased across sessions was similar for both groups of females.

### Switching Discrimination Task

The switching discrimination task (SDT) assessed discrimination learning. Figure 2 shows the mean discrimination index (DI) for all ICSI and CTL mice (top) and for ICSI/CTL males and females (bottom) in the SDT procedure. DI increased for all mice across the 20 training sessions. While both groups gradually made fewer unrewarded responses during this training, the top panel shows that the CTL mice made a greater proportion of rewarded responses from the third session onward and reached a substantially higher DI by the final session (0.68, +/- 0.02 SEM for CTL compared to 0.58, +/- 0.02 SEM for ICSI). The graphs in the bottom panels of Figure 2 show that both male and female CTL mice often had higher DIs than their ICSI counterparts, but the difference between CTL and ICSI discrimination performances was more pronounced and consistent for males.

**Figure 2.**
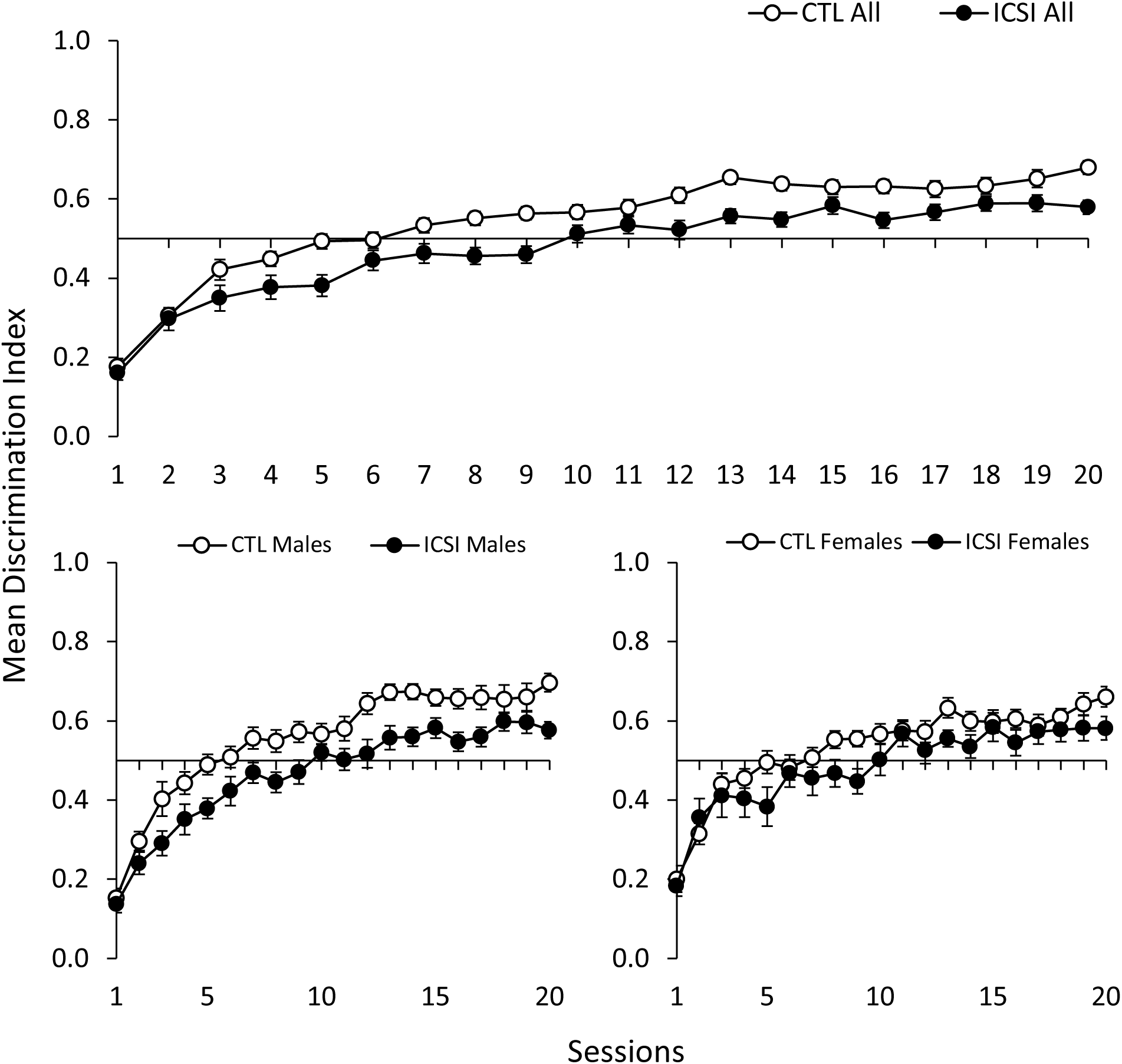
Mean discrimination index scores (+/- SEM) during switching discrimination task (SDT) sessions for all ICSI and CTL mice (top) and separated by ICSI and CTL males and females (bottom).

Statistical analysis for the comparison between all ICSI and CTL mice found a large main effect for session (F_19, 1349_ = 100.63, p < 0.001, ηp^2^ = 0.59) and a main effect for group (F_1, 71_ = 11.77, p = 0.001, ηp^2^ = 0.14), but no effect for the group x session interaction. Similarly, the comparison between ICSI and CTL males found significant main effects for session (F_19, 665_ = 71.16, p < 0.001, ηp^2^ = 0.67) and group (F_1, 35_ = 11.10, p = 0.02, ηp^2^ = 0.24) but no group x session interaction. The comparison between ICSI and CTL females found a significant main effect for session (F_19, 646_ = 35.96, p < 0.001, ηp^2^ = 0.51) but no main effect for group or the group x session interaction.

Taken together, CTL mice exhibited better discrimination learning, and this difference was more pronounced between CTL and ICSI males than it was between the female groups. Despite this, statistical analyses did not reveal significant group x session interactions for any of the comparisons, including the comparison between CTL and ICSI males. Overall, it appeared that the rate of improvement in DI scores across sessions was similar for the two groups, but the CTL mice nevertheless had consistently higher DI scores.

### Delayed Non-Matching-to-Position Memory Task

This procedure assessed working and reference memory. Figure 3 shows the mean proportion of correct/rewarded trials in DNMTP recognition memory sessions for all ICSI and CTL (top panel) and for ICSI/CTL males and females (bottom panels). The top panel shows that while the proportion of correct responses made by both groups increased across sessions, the CTL mice consistently made more correct responses from the seventh session onward. As in the previous SDT procedure, there appeared to be a larger difference in performance between males than females. CTL males made a greater proportion of correct responses than ICSI males in every session except the second. On the other hand, CTL and ICSI females made approximately the same proportion of correct responses until the 11^th^ session, after which CTL females made slightly more correct responses in most sessions.

**Figure 3.**
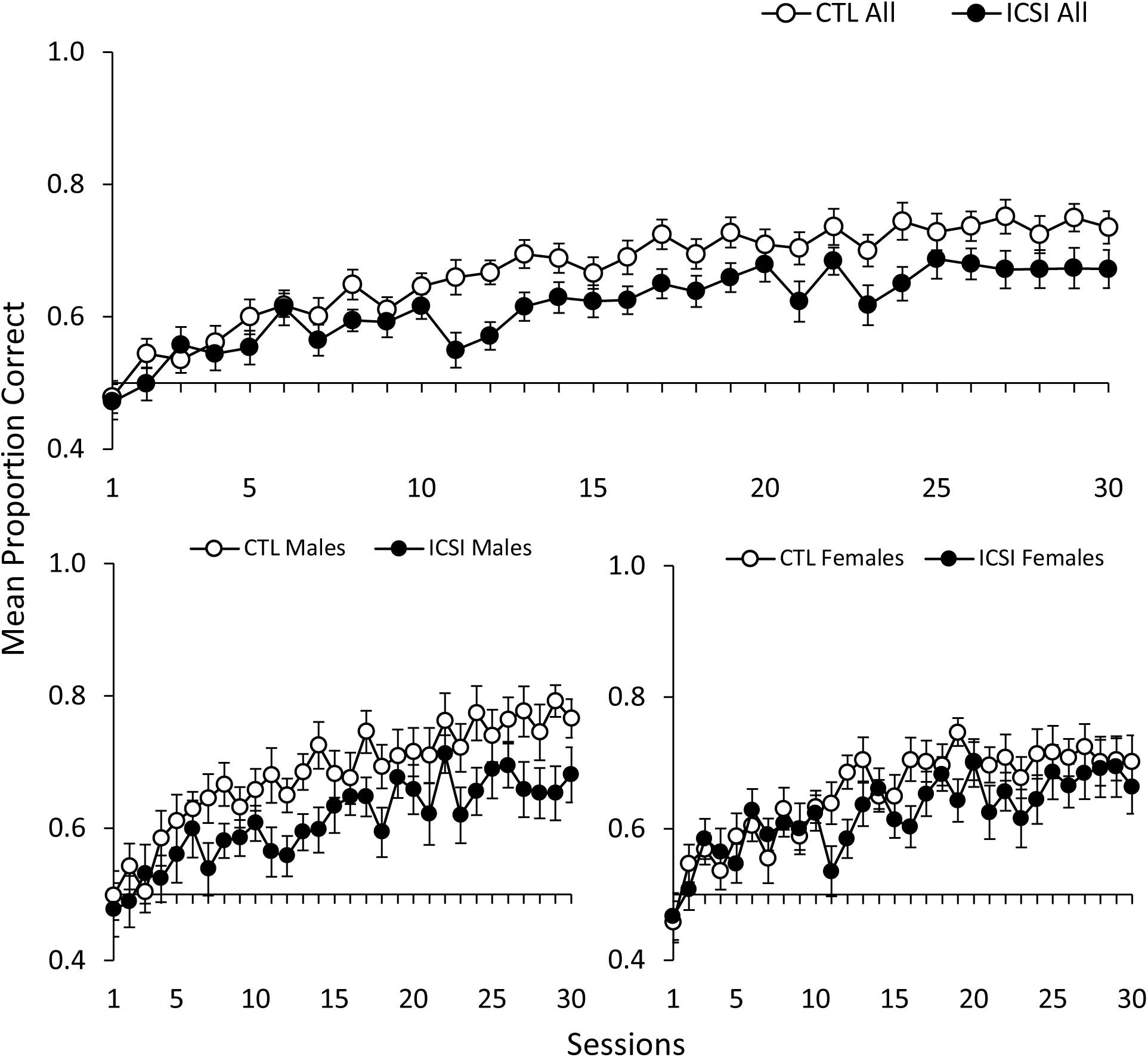
Mean proportion of correct trials (+/- SEM) in delayed-non-matching-to-position (DNMTP) sessions for all ICSI and CTL mice (top) and separated by ICSI and CTL males and females (bottom).

The mixed repeated measures ANOVA comparing all ICSI to CTL found significant main effects for session (F_29, 2030_ = 19.72, p < 0.001, ηp^2^ = 0.22) and group (F_1, 71_ = 7.67, p = 0.007, ηp^2^ = 0.10) but not for the group x session interaction. The comparison between CTL and ICSI males similarly found significant effects for session (F_29, 986_ = 10.80, p < 0.001, ηp^2^ = 0.24) and group (F_1, 71_ = 6.44, p = 0.02, ηp^2^ = 0.16) but not for the group x session interaction. For the comparison between CTL and ICSI females, there was an effect for session (F_29, 986_ = 9.59, p < 0.001, ηp^2^ = 0.22) but not for group or the group x session interaction.

The results of the DNMTP memory procedure were similar to those obtained in the preceding SDT. Specifically, CTL mice performed better than ICSI, and the difference between CTL and ICSI males was more pronounced than the difference between CTL and ICSI females. Statistical tests again found significant main effects for the group factor in the comparison between all ICSI/CTL and between male ICSI/CTL, but there was not a significant group x session interaction. Thus, it appeared that ICSI and CTL performance improved at approximately the same rate across training sessions, but the CTL mice (especially the males) consistently made a greater proportion of correct responses than their ICSI counterparts.

### DNMTP Retention Checks

Retention of the DNMTP performance was assessed with three retention check sessions. Figure 4 shows the mean proportion of correct responses in the three DNMTP retention checks for all ICSI and CTL mice (top) and separated by males and females (bottom). For reference, the first (leftmost) data point on these figures represents the mean proportion correct for each group in the last five DNMTP training sessions (i.e., sessions 25-30). The top panel shows that the mean proportion correct decreased slightly for both groups in the first (2-day) retention check relative to the last five sessions of DNMTP training. For CTL mice, the mean proportion correct continued to decrease slightly across the remaining two retention checks while the ICSI mice’ performance remained at approximately the same level. The two groups’ performances were equal in the final 10-day retention check. The bottom panels of Figure 4 shows that the proportion correct decreased monotonically for CTL males and females across the retention checks. For ICSI males, the proportion correct in the 5-day test increased slightly relative to the 2-day test but decreased to approximately the same level as the CTL males in the 10-day test. For ICSI females, proportion decreased across the 2- and 5-day tests but increased slightly in the final test.

**Figure 4.**
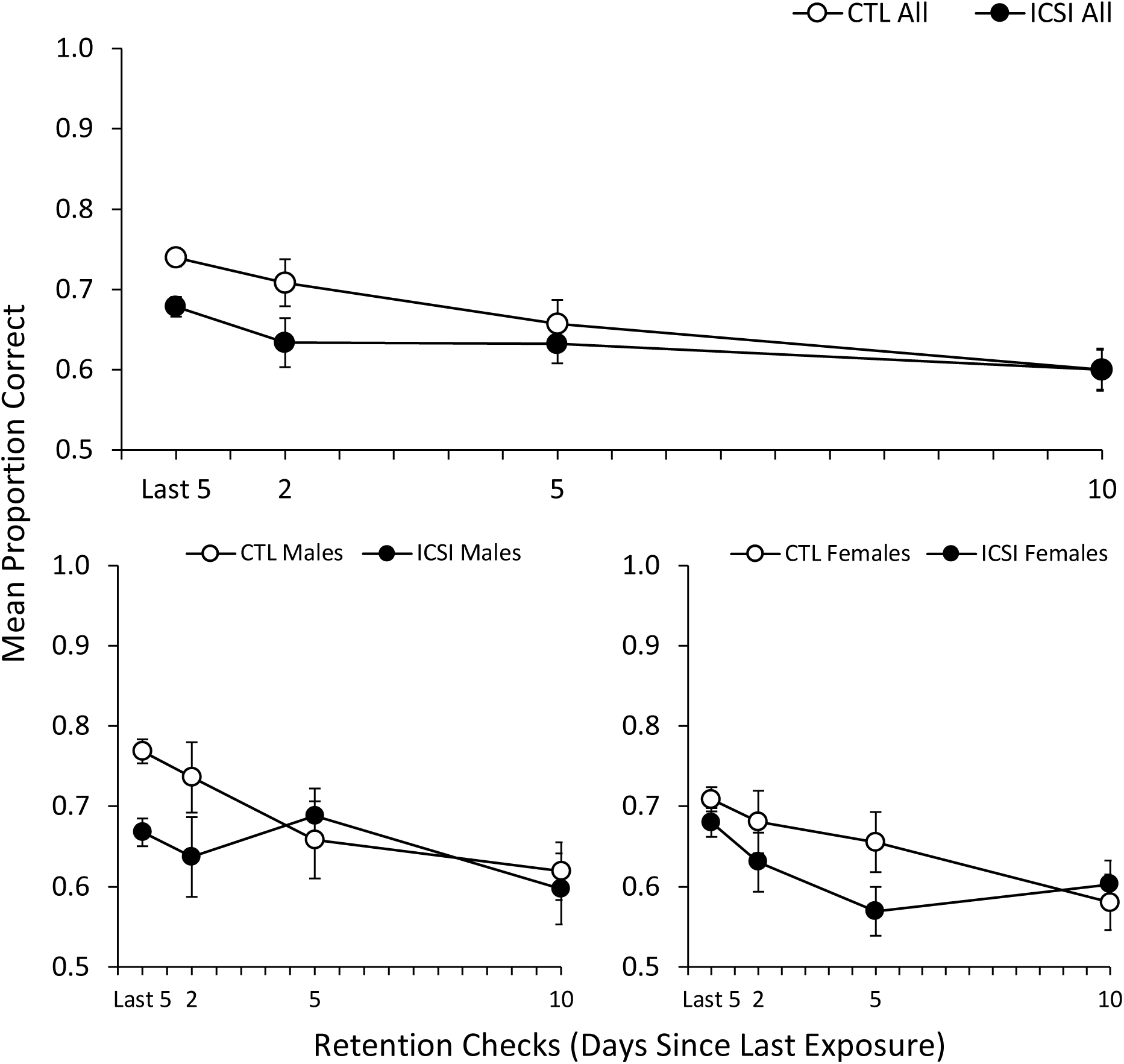
Mean proportion of correct trials (+/- SEM) in DNMTP retention checks for all ICSI and CTL mice (top) and separated by ICSI and CTL males and females (bottom). The leftmost data point represents the mean proportion correct by each group in the last five DNMTP training sessions.

**Figure 5.**
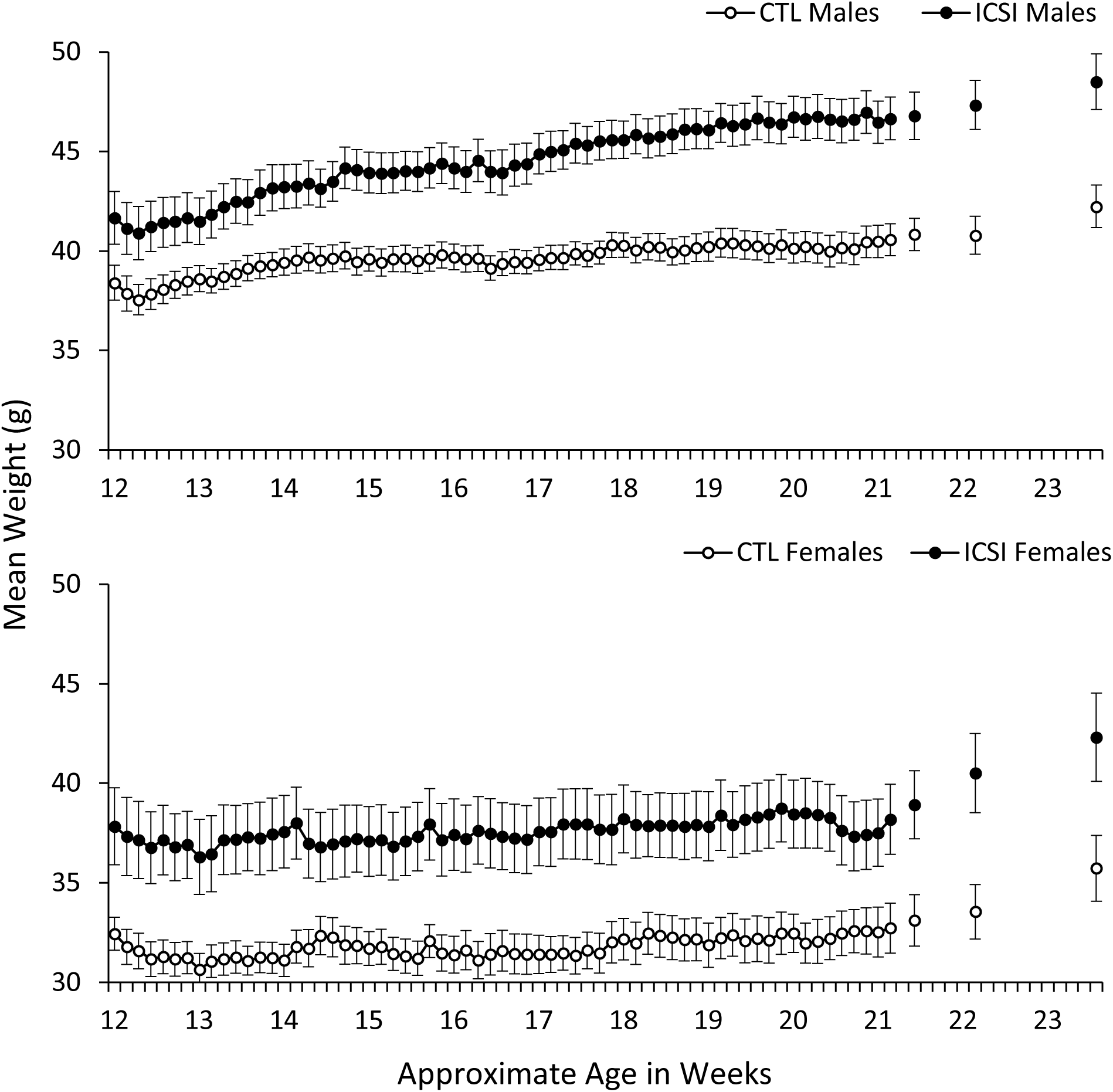
Mean daily weights in grams (+/- SEM) for ICSI and CTL males (top) and females (bottom) from the first session of magazine training to the final retention check. The x-axis shows the approximate ages of the mice in weeks. Weights were taken daily prior to sessions following a 14-h period of food deprivation. The gaps in the data series during weeks 21-23 were days between retention checks where mice were not weighed and had continuous free access to food.

The mixed ANOVA comparing all ICSI and CTL mice found a significant effect for session (i.e., significant decreases in proportion correct across the three retention checks; F_2, 140_ = 6.17, p = 0.003, ηp^2^ = 0.08) but no effects for group or group x session interaction. The comparisons between ICSI and CTL males and females likewise found significant effects for session for both (F_2, 68_ = 3.17, p = 0.05, ηp^2^ = 0.09 for males and F_2, 68_ = 3.65, p = 0.03, ηp^2^ = 0.10 for females), but found no effects for group or group x session interaction for either. Thus, while there was a general decrease in proportion correct across the three retention checks, there was no significant difference between the groups in the rate at which this decrease occurred.

### Follow-Up Assessments with Aged Mice

Follow-up assessments were conducted with aged mice to evaluate long-term retention and reacquisition of learned performances. Figure 6 shows the results for the SDT and DNMTP memory re-training sessions. As can be seen in the left panel, the mean DI for both groups improved slightly across the 15 SDT sessions and there was a significant effect for session (F_14, 434_ = 5.96, p < 0.001, ηp^2^ = 0.16). As during the initial SDT training, CTL mice showed better discrimination performances on average, but there was no significant effect for either group or group x session interaction. Discrimination improved for both groups across re-training, but neither group achieved the same level of performance as they had after the initial 15 SDT training sessions (cf., Figure 2).

**Figure 6.**
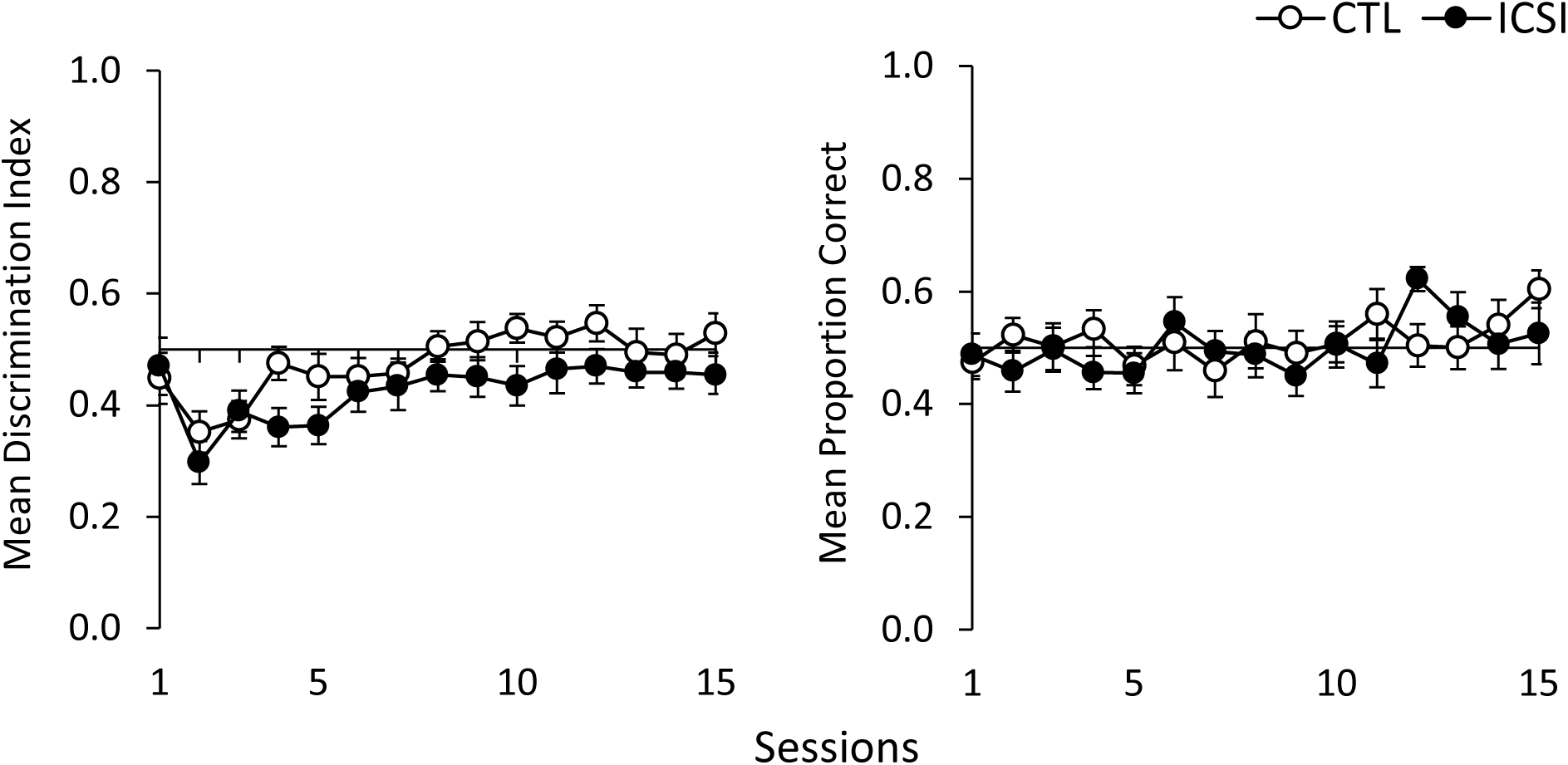
Mean discrimination index during SDT (left) and mean proportion of correct trials during DNMTP (right) follow-up assessments conducted when mice were 52-56 weeks of age. Error bars represent +/- SEM.

The right panel of Figure 6 shows performance in the DNMTP memory reassessments. The aged mice were unsuccessful in re-learning this performance after 15 sessions. Neither group approached the levels obtained after the 15 initial DNMTP sessions (Figure 3), and there was no discernible improvement beyond chance responding. There was no significant effect for session, group, or group x session interaction.

### Body Weight

As noted above, mice were weighed immediately prior to all sessions following a 14-h period of food deprivation. Figure 7 shows the mean daily weights of the ICSI and CTL males (top) and females (bottom) from the first session of magazine training to the final DNMTP retention check. The gaps in the data series during weeks 21-23 were days between retention checks where mice were not weighed and had continuous free access to food. Across the experiment, ICSI males and females both consistently weighed more than their CTL counterparts. Both ICSI and CTL males gained weight across the experiment, but ICSI males gained weight at a greater rate than CTL males. Compared to the males, the females gained relatively little weight across the experiment. However, both ICSI and CTL females gained a larger proportion of weight during the retention checks when they had longer periods of access to food. From the last DNMTP session to the final retention check, the weights for ICSI and CTL males increased by 3.94% and 3.42%, respectively. In comparison, ICSI female weights increased by 10.79% and CTL female weights increased by 7.88% during the same period.

**Figure 7.**
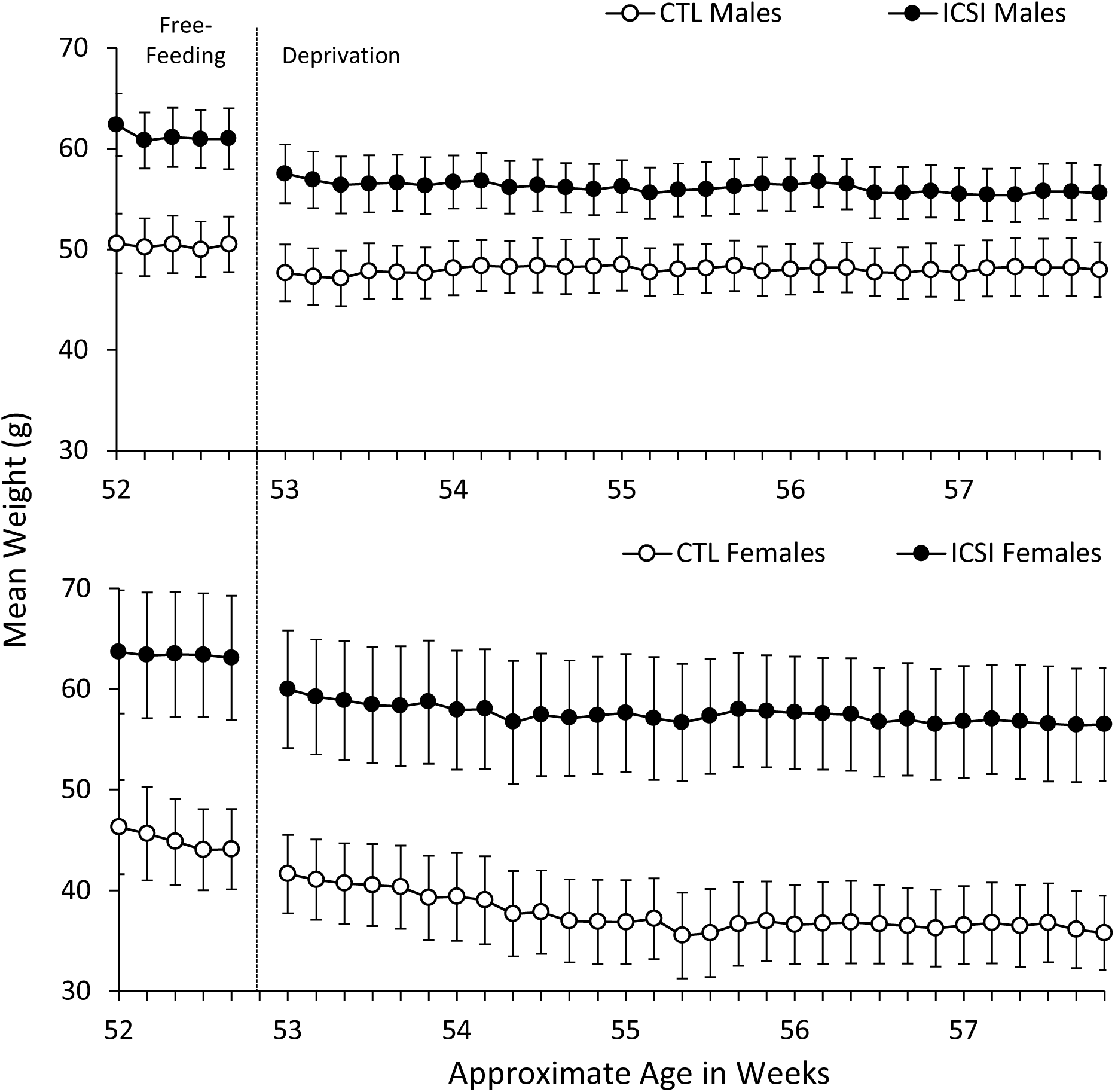
Mean daily weights in grams (+/- SEM) for ICSI and CTL males (top) and females (bottom) five days prior to and during the follow-up assessments. Weights to the left of the phase change line were taken daily while mice had free access to food. Weights to the right of the line were taken daily prior to sessions following a 14-h period of food deprivation.

Figure 7 displays the mean weights of the mice for five days prior to and during the reassessment training sessions starting at 52 weeks of age. All mice had *ad libitum* access to food from the end of the learning and memory initial assessments (when they were approximately six months of age) to the time of the re-training, when the food deprivation regimen was reinstated. At the first weighing after six months of free-feeding, ICSI males weighed an average of 62.4 g (+/- 3.10 SEM) compared to 50.6 g (+/- 2.96 SEM) for CTL males. ICSI females likewise weighed substantially more than their female CTL counterparts (63.7 g +/- 6.25 SEM for ICSI compared to 46.3 g +/- 4.67 for CTL).

The reinstatement of the food deprivation schedule produced an immediate reduction in weights of the males, but weights stayed largely the same until the end of the reassessments 30 days later. For females, the food deprivation schedule resulted in progressively lower weights across this same time, and this was more pronounced for the CTL females.

## Discussion

We subjected ICSI and CTL mice to a series of operant learning procedures to assess acquisition, discrimination learning, and memory. The inclusion of both males and females allowed for global comparisons between ICSI and CTL mice as well as for same-sex comparisons between the groups. Overall, CTL mice were found to outperform their ICSI counterparts in all but one of the learning and memory tasks we employed during their initial training, and the differences were largely due to sex-specific differences in performance in the tasks. Specifically, CTL females performed better during acquisition learning than ICSI females, but there was no difference in acquisition between ICSI and CTL males. In the SDT and DNMTP procedures, CTL males exhibited superior discrimination learning and memory compared to their ICSI counterparts, but there was not a statistically significant difference between ICSI and CTL females in these tasks. There were no apparent differences between the groups in the DNMTP retention checks designed to assess longer-term memory. Both groups showed significant decrements in performance in SDT and DNMTP re-training sessions conducted at 52 weeks of age.

While CTL mice exhibited superior performance in all procedures except the DNMTP retention checks during initial training, it is interesting to note that statistical analyses revealed significant group effects but no significant effects for group x session interactions. This means that the extent to which performance increased across training sessions was roughly equivalent for ICSI and CTL in the procedures employed here. Despite similar changes in behavior across repeated exposures to the learning and memory assessments, CTL mice consistently performed at a higher level. At this point it is unclear why this was the case. Further research investigating basic learning processes with these mice will be required to explain this difference.

A notable auxiliary finding was the relatively large and consistent difference in weights between ICSI and CTL mice. ICSI males and females both weighed more than their CTL counterparts both during initial training when mice were three to six months of age and when mice were over a year old. Other studies have similarly reported higher weights at birth for ICSI B6C3F1 males and females relative to CTL (Scott et al., 2010) as well as significantly higher weights for ICSI CD-1 females relative to CTL females from approximately 15 weeks of age (Fernández-Gonzalez et al., 2008). These data suggest that further investigations into potential metabolic differences between ICSI and CTL mice may be warranted.

There were limitations of the study that must be acknowledged. First, the study was not blinded: the technicians who handled the mice before and after their daily sessions were aware of the groups to which they belonged. Although the training sessions (including the recording of data) were entirely automated and the technicians’ interactions with the mice were limited to weighing and transporting to and from the experimental chamber in handling tubes, blinding would add an additional level of rigor and control for any inadvertent differences in how mice were handled. A second limitation is that the procedures were conducted in succession, meaning that each individual assessment occurred when the mice were at a single age. It may be the case that comparing acquisition, discrimination, or memory between ICSI and CTL mice at different points in the developmental timeline may yield different results. As a proof of concept study, we aimed to show the potential effects of the overall ICSI procedure on the health of offspring; thus, we did not distinguish multiple factors involved in ICSI, e.g., superovulation protocol, sperm preparation protocol, culture conditions, injection conditions, stages for embryo transfer, and the age of surrogate mothers. These variables may be worth testing in future studies.

Despite these limitations, the present study strongly suggests that studying learning and memory in animal models has the potential to shed light on outcomes of ICSI at the level of cognitive function. Our data open up a number of avenues for further investigation. In this study, we investigated operant learning and memory using only reinforcement procedures in which sugar pellets served as the reward. Studies have shown that mouse models that exhibit learning deficits relative to control mice in one type of operant procedure may exhibit superior performance in a different operant learning paradigm (Lewon et al., 2017). It is therefore necessary to expose mouse models to as many types of learning situations as possible to obtain the fullest picture of cognitive function. Operant learning assessments are diverse and include procedures that use other types of rewards under different schedules of reinforcement, different types of spatial and multisensory discrimination and memory tasks, escape/avoidance learning tasks, and procedures that provide measures of sensitivity to stress-inducing aversive events. In addition to operant learning procedures, future studies may also examine more basic processes such as nonassociative and Pavlovian learning. One benefit of the modular experimental chambers such as those used in this experiment is that a single apparatus may be readily modified to accommodate all of these types of assessments. As there appeared to be sex-specific differences in learning and memory in this experiment and studies have similarly found evidence of sexual dimorphism in other measures of ICSI outcomes (Esteves et al, 2018; Fernández-Gonzalez et al., 2008), this research should include assessments of both males and females (Shansky, 2019).

In addition to studying ICSI mice with other types of learning procedures, future research may also examine how variables related to the ICSI procedure itself may affect learning and memory. Some studies have found that ART is associated with an increased occurrence of epimutations and imprinting disorders (de Waal, et al., 2012; Lazaraviciute et al., 2014; Pinborg, 2016), and it is known that ARTs may induce embryonic stress responses that alter gene expression and exert a number of other epigenetic effects during early development (Ramos-Ibeas et al., 2018; Szöke et al., 2018). Laboratory procedures related to ICSI (e.g., sperm extraction and selection methods, sample handling, egg retrieval and culture, etc.) may further contribute to the likelihood of epigenetic alterations (Esteves et al., 2018; Ghosh et al., 2017; Palermo et al., 2017). Environmental events occurring during lifetime of individuals are known to produce modifications in gene expression that affect neurodevelopment and psychological function across the lifespan (Grigorenko et al., 2016; Guan et al., 2015), and there is evidence that some of these modifications may be inherited by offspring (Babenko et al., 2015; Chen et al., 2016; Nestler, 2016; Jablonka & Raz, 2009). For all of these reasons, future research should investigate how the ICSI procedure and the epigenetic factors associated with it affect cognitive function, ideally across multiple generations.

It is premature to speculate as to the implications of these results to cognitive function and the psychological development of ICSI humans. Although ICSI mice exhibited certain learning and memory deficits relative to CTL mice in the testing we employed, cognitive deficits should not be assumed to be invariably associated with ICSI in humans. There are several reasons for this. First, as noted above, the assessments conducted here represent a small portion of the procedures available for investigating learning and memory, and a wider range of these will be needed to more fully characterize cognitive function in ICSI mice. Second, human learning environments differ in important ways from mice (Hayes & Delgado, 2007), and families of ICSI children vary widely in terms of socioeconomic status, education, and access to medical and educational resources for their children. The deficits observed in ICSI mice in this study may therefore prove to be clinically insignificant in certain social environments.

Finally, and perhaps most importantly, cognitive function must be seen as the product of a complex set of interactions between individuals and their environments throughout the lifespan. During development, environmental factors interact with genetic materials to determine the physiological phenotypes of whole individuals. These individuals then interact with their physical and social environments, which shape their behavior across time through nonassociative, Pavlovian, and operant learning processes. Different learning environments will inevitably impart different repertoires, and the physiological characteristics of individuals (e.g., brain function, metabolism, sensory abilities, etc.) determine their capacity for learning from particular types of environmental contingencies. Physiological characteristics that provide advantages for learning in certain environments may prove to be detrimental in others (Lewon et al., 2017). For these reasons, studying the relationship between ICSI and cognitive function is a truly interdisciplinary endeavor that does not fall solely within the domain of either genetics or psychology (Hayes & Fryling, 2009). Genetic and epigenetic analyses by themselves cannot explain cognitive development in a directly causal manner, as this depends in large part upon the types of interactions individuals have with their environments. Similarly, analyses at the psychological level alone cannot explain differences in learning capacities related to genetic characteristics. Further interdisciplinary research on basic learning processes with mouse models has the potential to enhance our understanding of these interactions as they relate to ICSI and other ARTs. This research will require close coordination between investigators at both the genetic and psychological levels of analysis.

## Acknowledgements

The authors would like to thank John F. Gray, Cole Gansberg, Taylor Chase, Laura Cohen, Kristen Green, Osmar Lopez, Elisabeth Mclean, Haley Mizell, Caitlyn Peal, Keenan Raquel, Tori Sandoval, Emily Spurlock, Melanie Stites, and Jamiika Thomas for technical assistance.

## Role of Authors

ML, YW, CP, and MP contributed to the initial draft of the manuscript, and subsequent edits were made by all authors. ML, YW, CP, HZ, LH, and WY contributed to the conception and design of the study. The ICSI procedure and breeding were conducted by YW and HZ. Learning and memory assessments and data analysis were conducted by ML, CP, and MP.

